# Transcriptional Biomarker Discovery Towards Building A Load Stress Reporting System for Engineered *Escherichia coli* Strains

**DOI:** 10.1101/2023.03.28.534627

**Authors:** Yiming Huang, Anil Wipat, Jaume Bacardit

## Abstract

Foreign proteins are produced by inserting synthetic constructs into host bacteria in biotechnology applications. This process can cause resource competition between synthetic circuits and host cells, placing a metabolic burden on the host cells which may result load stress and detrimental physiological changes. Consequently, the host bacteria can experience slow growth, while the synthetic system may suffer from suboptimal function and reduced productivity. To address this issue, we developed machine learning strategies to select a minimal number of genes that could serve as biomarkers for the design of load stress reporters. We identified pairs of biomarkers that showed discriminative capacity to detect the load stress states induced in 41 engineered *E. coli* strains. These biomarker genes are mainly involved in Envelope stress response, Ion transport, Energy production and conversion.

## 1 Introduction

In the biotechnology industry, prokaryotic expression systems are utilised to yield valuable products such as enzymes, chemicals, pharmaceuticals and biofuels. *Escherichia coli* remains the preferred host strain to be engineered for the generation of diverse proteins [1, 2]. Most of these proteins are not naturally found in *E. coli*, thus synthetic constructs are designed and inserted to enable the introduction of foreign genes and proteins in the host strain [3]. Unfortunately, the expression of the synthetic constructs can excessively consume the cellular resources [4, 5] and impose an unnatural metabolic burden on the bacteria [6]. The load stress resulting from this metabolic burden can trigger detrimental physiological adaptations that potentially harm not only the growth of the host bacteria [7, 8] but also the performance of the synthetic system as a whole [9, 10, 11]. Therefore, understanding the host strain’s reaction to metabolic load and monitoring the cellular load stress are essential for successful synthetic construct expression.

Bacteria grow in environments that are constantly changing, often subjecting them to periods of multifarious stress conditions. As a result, they have evolved mechanisms to induce gene expression, metabolic and physiological changes that can help mitigating against the damage resulting from these stresses[12, 13]. Transcriptomics technologies, such as RNA-seq [14], have revolutionised our ability to study bacterial transcriptomes in different conditions, allowing us to investigate how they shift in response to changing environments and infer the gene regulatory parts involved in stress responses [15, 16]. Focusing on the load stress that the engineered strains can undergo while heterologous proteins expression, some studies have explored the transcriptional changes in host cells and identified key biomarker genes that react to load stress [17, 18]. Additionally, feedback control systems incorporating these reporting genes or promoters have been developed to recognise load stress states and adjust synthetic construct expression [19, 20, 21].

These previous load stress studies, however, have looked at only a few synthetic constructs and a small number of foreign proteins, leading to restricted understanding of the host response to load stress. The resulted load stress reporting systems in these studies may therefore fail to recognise the cellular states induced by various unseen foreign protein expression. Meanwhile, these prior research have been also limited on transcriptional changes to load stress, missing comparison between load stress and other stress states (e.g., heat, acid, nutrients scarcity) in the host cells which are also common in industrial production [22, 23]. The biomarker genes identified in these cases are thus not unique to load stress and can mistakenly detect other stress responses that cause similar expression shifts in these load stress reporting genes.

This study aimed to develop machine learning methods for pinpointing a few key genes in *E. coli* that can discriminate load stress state induced by expressing a larger set of heterologous genes, with respect to a wide range of other cellular states presented by growing in various environments. We hypothesised that a minimal number of genes can be identified to indicate load stress by mining a large-scale transcriptome that contains both samples induced with many different heterologous genes and samples grown in assorted conditions. A recent compendium of *E. coli* RNA-seq data Precise2 [24] is such a transcriptomics dataset, on which we studied the transcriptional response to load stress and developed an ensemble of feature selection models to optimally decide the number of genes required to sense load stress. We highlighted the biomarker genes with great prediction performance and discriminative power. The identified biomarker genes can be harnessed to improve the performance of the burden feedback systems to monitor and relieve the load stress states elicited by producing a wide range of foreign proteins in *E*.*coli* cells.

## 2 Methods and data

### 2.1 RNA-seq data acquisition and normalisation

PRECISE 2.0 [24] is a RNA-seq compendium, available at https://github.com/SBRG/precise2, for *Escherichia coli* K-12. This dataset is suitable for exploring the load stress transcriptional response because it collects transcriptomes from a large-scale heterologous gene expression experiment and many other experiments applying gene and environmental perturbations while growing the bacteria. To identify transcriptional biomarkers for load stress state in *E. coli*, we analysed 251 gene expression profiles from this dataset. These include a) 169 samples of MG1655 strain grown with various environmental perturbations such as alternative carbon sources, alternative nitrogen sources, nutrient limitation, antibiotic drugs, anaerobic condition and other stimulus (e.g., acid, oxidative drugs, ethanol, NaCl) and b) 82 samples of 41 engineered strains which respectively expressed a heterologous gene or had an empty plasmid inserted.

Lamoureux et al. processed the raw data using a Nextflow pipeline (https://github.com/avsastry/modulome-workflow) designed for microbial RNA-seq datasets and reported read counts and log-transformed Transcripts per Million (log-TPM) after quality control. We used the gene expression read counts for the differential expression analysis. For biomarker discovery study with feature selection methods, we further normalised the log-TPM to minimise the between-experiments noises by subtracting study-specific reference expression quantities. The study-specific reference expression quantities were calculated as the average expression levels of control samples from the same study wherever possible. In the studies where the control experiments were not conducted, the reference expression quantities were computed from the samples grown with least environmental perturbation factors due to different base medium or carbon sources. Please find the reference sample list and normalised gene expression quantities in Supplementary Table 1.

### 2.2 Identifying differentially expressed genes and overrepresented Gene Ontology terms

The R package DESeq2 [25]was applied to identify differentially expressed genes (DEGs) on 251 gene expression profiles collected in various conditions. We compared pairs of sample groups defined by 53 environmental perturbation factors (Supplementary Table 2a), including heterologous gene expression conditions that can induce load stress states, treatments of different stimuli, alternative nutrients or medium bases, etc. The significance threshold of DEGs was set as *log*-*fold*-*change >* 1 and *p*-*value <* 0.05 (obtained by Wald statistical test with multiple testing corrected by the Benjamini and Hochberg method).

Gene Ontology (GO) enrichment analyses was performed to study the overrepresented functions in these DEGs. Fisher exact test with Bonferroni correction was used to select the GO items with false discovery rate less than 0.01.

### 2.3 Regulon and imodulon activity analysis

A regulon is a group of genes or operons that are turned on or off in response to the same signal by the same regulatory protein. The regulons used in this study were taken from an *E. coli* transcriptional regulatory network which integrates RegulonDB v10.5 [26], Ecocyc [27] and a recent study about uncharacterised transcriptional factors [28]. This regulatory network consists of 371 regulons and 3407 targeted genes.

An imodulon, which was coined by Sastry, et al [29], is a group of genes representing an independently modulated signal that are likely controlled by the same or related regulators. Therefore, imodulons can be inferred from large scale transcriptomic data to suggest hypothetical regulatory mechanisms in addition to prior known regulons. Here we analysed the imodulons generated by [24] from PRECISE2 transcriptomics data for *E. coli*. The total of 218 imodulons are listed and characterised in an *E. coli* imodulon database https://imodulondb.org/search.html?organism=e_coli&dataset=precise2.

We summarised the activities, reflected in percentage index and intensity index, of regulons and imodulons underlying the transcriptional changes in 53 environmental perturbations. The percentage index of a given regulon or imodulon for the studied environmental perturbation was computed as the proportion of DEGs that are matched in this regulon or imodulon. The intensity index of a given regulon was computed as the average fold changes of the matched genes. The intensity index of a given imodulon was calculated as the fold changes of the imodulon expression—each iModulon has a weight for each gene.

### 2.4 Feature selection methods

To identify a biomarker panel consisting of a few genes whose expression patterns can indicate load stress states, a feature selection process was performed to select a reduced set of features capable of classifying load stress samples from the samples grown in other conditions. We applied four feature selection methods: RFE+RF, RFE+SVM, RGIFE+RF, RGIFE+SVM, which are two feature elimination strategies Recursive Feature Elimination (RFE) [30] and Rank-Guided Iterative Feature Elimination (RGIFE) [31], in combination with a non-linear classifier Random Forest (RF) or a linear classifier Support vector machine (SVM).

RFE starts with an initial set of features and prunes a fixed number of least important features recursively until the desired number of features to select is eventually reached. The importance of each feature at each step is obtained based on the classifier model trained with the current set of features. Here we set the number of features to eliminate at each step as two.

RGIFE starts from a full feature set, iteratively drops a dynamic block of features only if their removal has no negative effect on the predictive capacity of the current classifier model. The predictive performance of the model is estimated from stratified 10-fold cross-validation. The order in which features are checked for removal is determined by the ranking of feature importance produced by the classifier as part of its training process. The block size of the features to be removed is set as 25% of the current feature size. If the trial to remove a block fails (because its removal led to a model of worse quality) the next block in order will be attempted. After five consecutive failed trials or all current features have been checked, the block size will be divided by 4 and the process will start again. Once the block size is reduced to 1, the algorithm will stop after five unsuccessful trials and will return the remaining features as the final panel, hence automatically deciding when to stop the iterative feature elimination process.

### 2.5 Determining the biomarker panel size and prioritising the biomarker genes

To determine the optimal number of genes required in a biomarker panel, we repeatedly ran the aforementioned four feature selection models to obtain sufficient biomarker panels that vary in size for predicative capacity comparison. The desired number of features (biomarker genes) in RFE was set as 2 to 6 and each configuration was run for 100 times. For RGIFE, we reran the model if the size of the final feature set does not fall between 2 to 6 until the experiments returned 100 biomarker panels for each panel size. We applied stratified 10-folds cross-validation to examine how the identified biomarker panels, as grouped by number of genes they consist, would perform in terms of f1-score.

To prioritise a few biomarker panels for further biological interpretation and evaluation, we assessed the identified biomarker panels using three criteria, i.e. f1-score, occurrence and margin. The occurrence was computed as the number of times a biomarker panel was selected by different models in multiple runs. The margin was computed as the minimal distance between load stress samples and other samples in the hyperspace of biomarker gene expression. As shown in equation 1, **K** denotes a set of genes in a given biomarker panel, while **X**_**I**,**K**_ and **X**_**J**,**K**_ are the expression levels for biomarker genes (**K**) in load stress samples (**I**) and other samples (**J**) respectively. The margin can be negative when the expression changes of all biomarker genes between sample i and sample j showed the opposite trends than the mean expression changes between load stress samples and other samples.

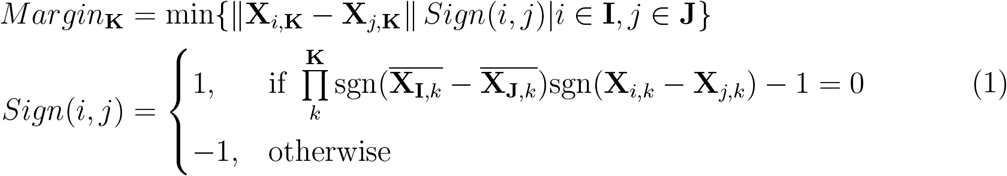

## 3 Results

### 3.1 Transcriptional response to load stress in *E. coli*

We compared the transcriptional profiles in heterologous gene expression samples, which were considered to induce load stress states in *E. coli*, with respect to wide type strain samples grown in control conditions. We detected 222 genes that presented significantly varied transcriptional levels as being responsive to load stress (Figure 1a). The gene ontology enrichment analysis revealed common biological processes underlying these differentially expressed genes, notably including the terms putatively related to excessive protein expression such as structural constituent of ribosome, translation, rRNA binding, cell envelope Sec protein transport complex, etc (Figure 1b).

**Figure 1:**
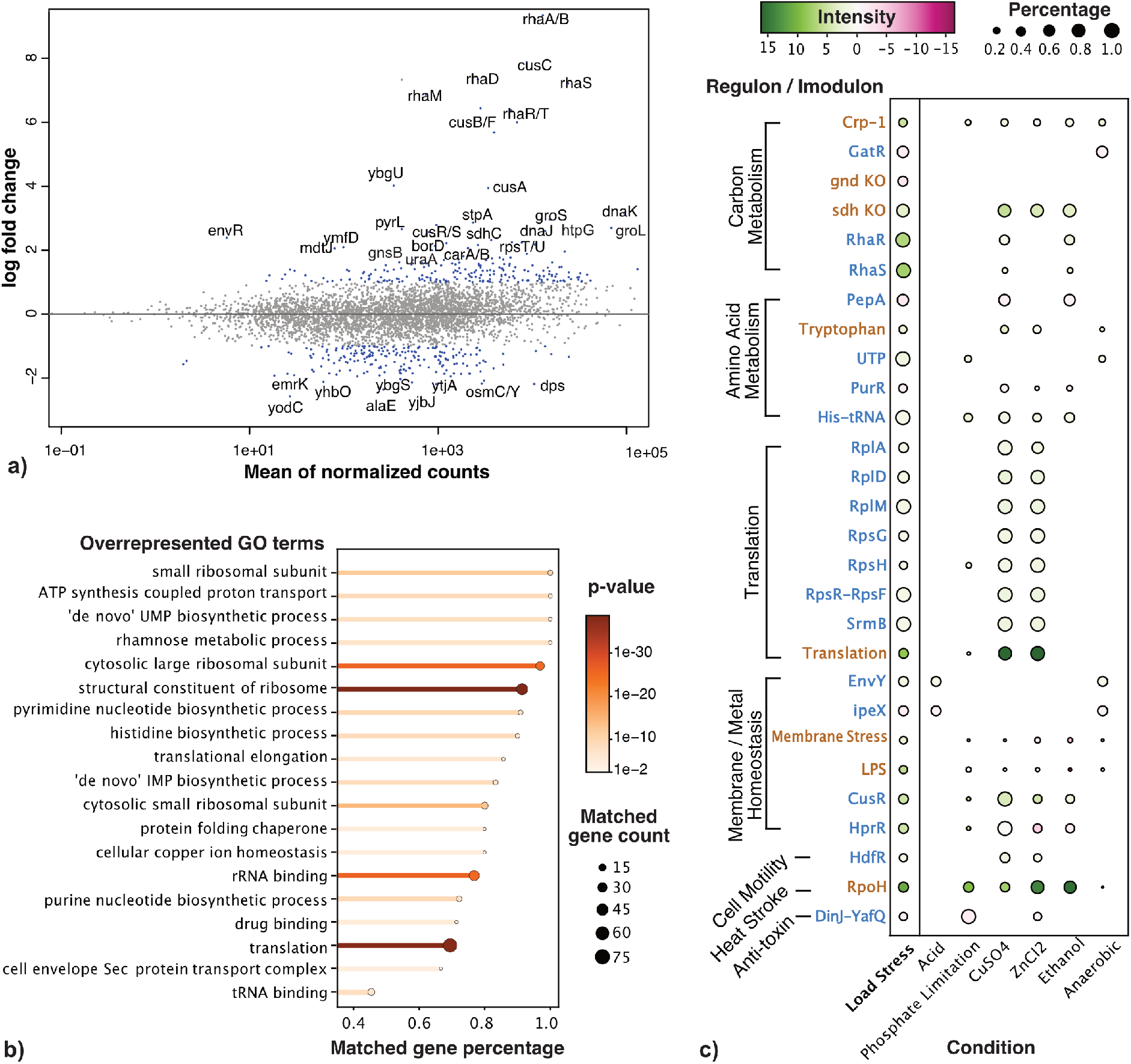
Transcriptional response to load stress in *E. coli*. **a)** The scatter plot shows the log fold change in gene counts between wide type strain samples and engineered strains, as against to the mean gene counts across all samples. The differentially expressed genes (DEGs) are highlighted in colour blue (p-value*<* 0.05, absolute log-fold-change*>* 1) with top DEGs labelled (absolute log fold change*>* 2). **b)** The overrepresented gene ontology (GO) terms are listed and ordered by “matched gene percentage”, i.e. the percentage of genes in a given GO term that present in the list of DEGs. The colour indicates the fisher exact test p-value and the end dot size indicates the “matched gene count”. **c)** The dot plot shows the activity level changes for a number of regulons and imodulons in response to several stress conditions as compared to load stress condition. The percentage index, reflected in dot size, indicates the percentage of genes associated with a given regulon or imodulon that are differential expressed in a given condition. The intensity index, reflected in dot colour, indicates the average fold change across genes matched in a given regulon or imodulon for a given condition. The regulators and imodulons with varied activity levels (percentage index*>* 0.3) for load stress condition are displayed, with regulons listed in colour blue and the imodulons in blown.

We further studied the transcriptional response to load stress by analysing the transcriptional regulator and imodulon activity changes in heterologous gene expression samples, as compared with the corresponding transcriptional alterations resulting from different environmental perturbations (Supplementary Table 2b-c). As shown in Figure 1c, the regulons and imodulons that were responsive in load stress condition mainly involve Carbon and Amino Acid Metabolism, Translation, Membrane or Metal Homeostasis, Cell Motility, Heat Stroke and Anti-toxin. Among which the regulators associated with Translation and Heat Stroke (such as RplADM, RpsFHR, RpoH) adjusted their activity levels in conditions treated with CuSO4 or ZnCl2, and the regulators associated with Membrane Homeostasis (EnvY, ipeX) adjusted their activity levels in acid stress condition and anaerobic condition, suggesting complex transcriptional alterations are triggered in load stress response.

We noted that the load stress samples show the overrepresented GO term “rhamnose metabolic process” as well as most significantly increased activities in regulators RhaS and RhaS. This effect is arguably due to the use of rhamnose as inducer in all heterologous gene expression samples, thus the relevant genes should be removed from the candidate biomarker genes for sensing the transcriptional state incurred by load stress.

### 3.2 Pairs of genes can serve as transcriptional biomarkers to sense load stress

We performed the feature selection process repeatedly on training folds of data with four different models, and estimated the overall performance by using the selected biomarker genes to predict the test folds of data. We obtained the cross-validation f1-scores for 2291 biomarker panels of size varying from 2 to 6 and by four models (Figure 2a). RFE-SVM and RGIFE-SVM embedded with SVM classifier performed better for selecting panel size larger than 2. RGIFESS-RF and RGIFESS-SVM adoptings dynamic feature elimination strategy performed better than RFE-RF and RFE-SVM in general.

**Figure 2:**
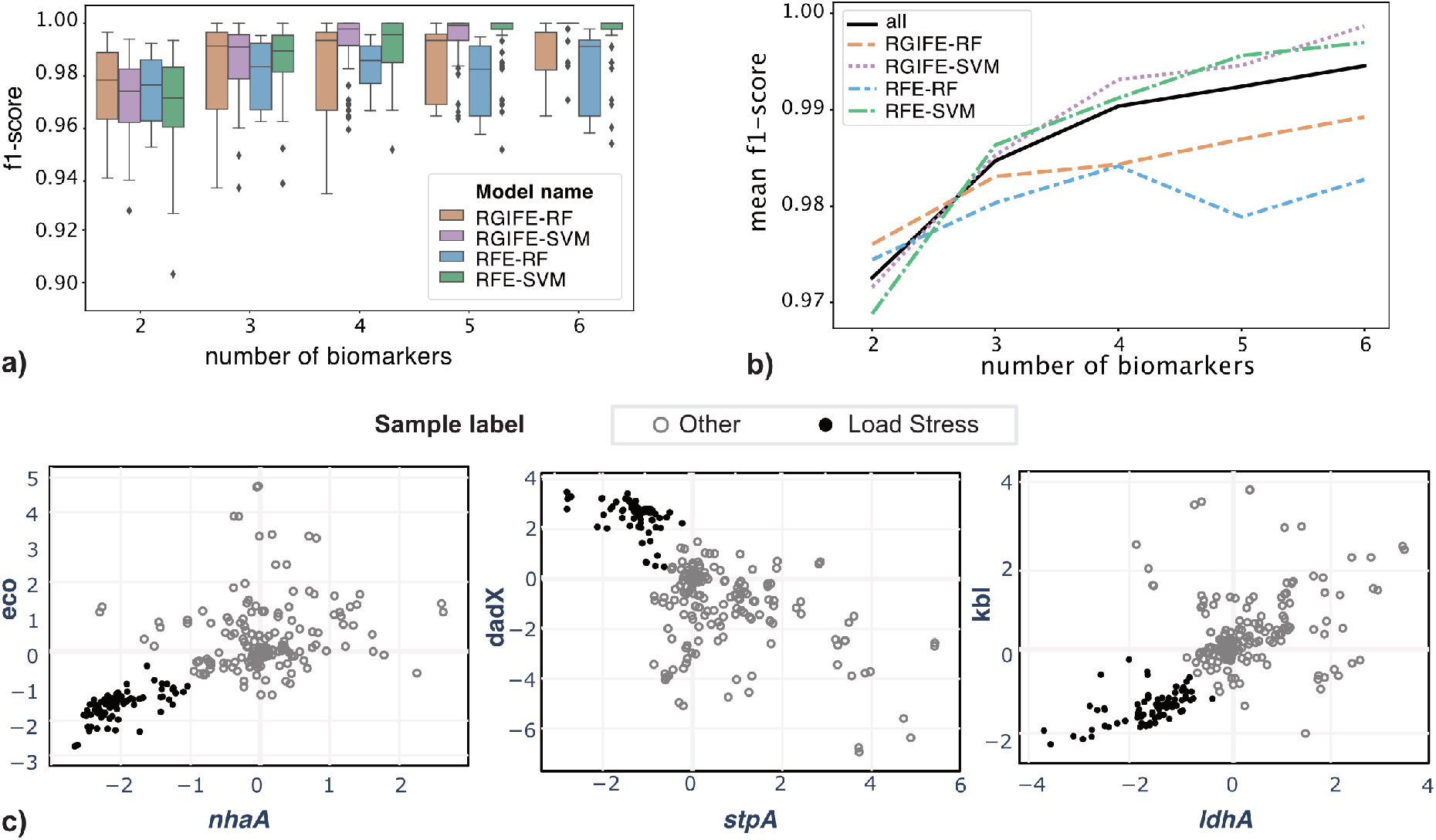
The performance of various biomarker panels to predict load stress in *E. coli*. **a)** The performance distribution across 100 repetitions of cross-validation tests for biomarker panels selected by four different models and of varying sizes (from 2 to 6). **b)** The mean performance for each of four models and all models combined. **c)** The gene expression levels of three pairs of identified biomarker genes are shown in the scatterplots. The black circle markers represent the samples from the load stress group while the grey open circle markers represent the other samples grown in various conditions.

The overall performance for all models except RFE-RF improved as the biomarker panel size increased (Figure 2b). However, building more biomarker genes in the load stress reporter system can add challenges and costs in the design of the system. We found that the biomarker panels consisting of two or three genes are able to discriminate load stress samples with median f1-score at 0.972 and 0.984, respectively.

We reported 13 biomarker panels consisting of two genes that achieved great predictive power (f1-score*>* 0.98, margin*>* 1) in Table 1, from which we highlighted in bold font a) a biomarker pair with the highest occurrence which was most stable across multiple runs of different biomarker identification methods, b) a biomarker pair with the highest f1-score which most accurately predicated load stress states, and c) a biomarker pair with the highest margin which showed the largest effect size to discriminate the load stress samples from other samples. The gene expression patterns of these biomarker pairs in load stress samples were distinct from the other samples (Figure 2c), and thus can be coupled to build synthetic systems capable of predicting load stress.

**Table 1:** The shortlist of biomarker pairs.

| Biomarker Genes | Model Names | Occurrence | f1-score | Margin |
| --- | --- | --- | --- | --- |
| ***nhaA;eco*** | RGIFE-RF;RFE-RF | 32 | 0.993 | **0.397** |
| *eco;queE* | RGIFE-RF;RFE-RF | 13 | 0.987 | 0.293 |
| *nhaA;yhcN* | RGIFE-SVM;RFE-SVM | 4 | 0.989 | 0.276 |
| *yodC;ygbA* | RGIFE-SVM;RFE-SVM | 9 | 0.984 | 0.216 |
| *gnsB;pabA* | RFE-SVM | 1 | 0.993 | 0.200 |
| ***ldhA;kbl*** | RGIFE-SVM | 10 | **0.997** | 0.195 |
| *cusC;stpA* | RGIFE-SVM | 6 | 0.994 | 0.194 |
| *nhaA;mdlA* | RGIFE-RF | 1 | 0.996 | 0.192 |
| ***dadX;stpA*** | RGIFE-RF;RFE-RF | **40** | 0.992 | 0.184 |
| *pqiA;eco* | RGIFE-RF | 12 | 0.991 | 0.165 |
| *nhaA;aaeA* | RGIFE-RF | 1 | 0.991 | 0.140 |
| *aaaD;ygbA* | RFE-SVM | 1 | 0.987 | 0.124 |
| *dadX;ldhA* | RGIFE-RF | 1 | 0.987 | 0.110 |

### 3.3 Biological characterisation of the biomarker genes

We analysed the functions of three highlighted biomarker pairs and found that at least one gene from each biomarker pair is involved in ion (e.g, Inorganic ion, Amino acid or Coenzyme) transport and metabolism (Table 2). To study the transcriptional factors that are directly or indirectly associated with our load stress biomarkers and the chains of transcriptional activities between our load stress biomarkers, we extracted a smallest component from *E. coli* Gene Regulatory Network that connects these six biomarker genes (Figure 3). In this sub gene regulatory network we noticed that the transcriptional factors Lrp, RpoD, ppGpp and h-NS, which regulate more than two biomarker genes, are generally related to Envelope stress response, Ion transport, transcription repression, Energy production and conversion.

**Table 2:** Gene annotations for three highlighted *E. coli* load stress biomarker pairs.

|  | Gene | Product | Function |
| --- | --- | --- | --- |
| Biomarker Pair 1 | *nhaA*<br>*eco* | Na(+):H(+) antiporter NhaA<br>serine protease inhibitor ecotin | Inorganic ion transport and metabolism<br>Cell wall/membrane/envelope biogenesis |
| Biomarker Pair 2 | *dadX*<br>*stpA* | alanine racemase 2<br>DNA-binding transcriptional repressor StpA | Amino acid transport and metabolism<br>Transcription |
| Biomarker Pair 3 | *kbl*<br>*ldhA* | 2-amino-3-ketobutyrate CoA ligase<br>D-lactate dehydrogenase | Coenzyme transport and metabolism<br>Energy production and conversion |

**Figure 3:**
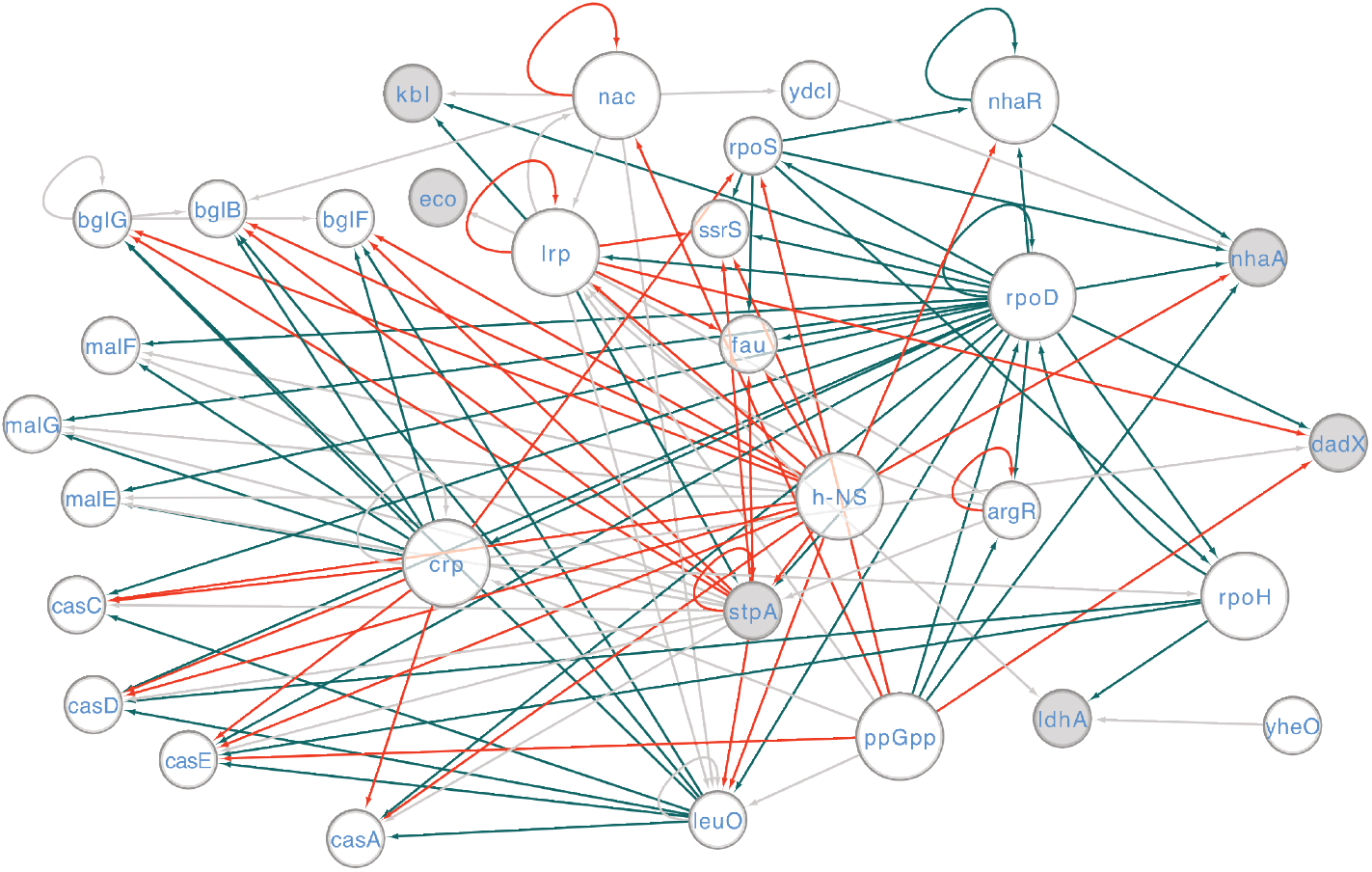
The subnet of *E. coli* transcriptional regulatory network related to load stress biomarker genes. The sub transcriptional regulatory network that contains the shortest path connecting genes in three selected biomarker pairs. The biomarker genes are highlighted in colour grey and the associated regulators are highlighted in larger node size.

To further characterise the transcriptional behaviours of the identified biomarker genes, we compared the expression levels of the biomarker genes in load stress samples with their expression in other samples grown in various conditions. As shown in Figure 4a, the relative expression levels reflect the transcriptional changes incurred by moving the samples to 23 treatment condition from the corresponding control conditions. We found that at least one gene in a biomarker pair presented the opposite trend or much larger scale of the same trend of transcriptional changes in response to load stress as against most of the other conditions. However, multiple individual biomarker genes saw similar transcriptional changes in some stimuli (such as Dibucaine, CuSO4 and Alkali) as in load stress, suggesting that common stress response mechanism may be shared.

**Figure 4:**
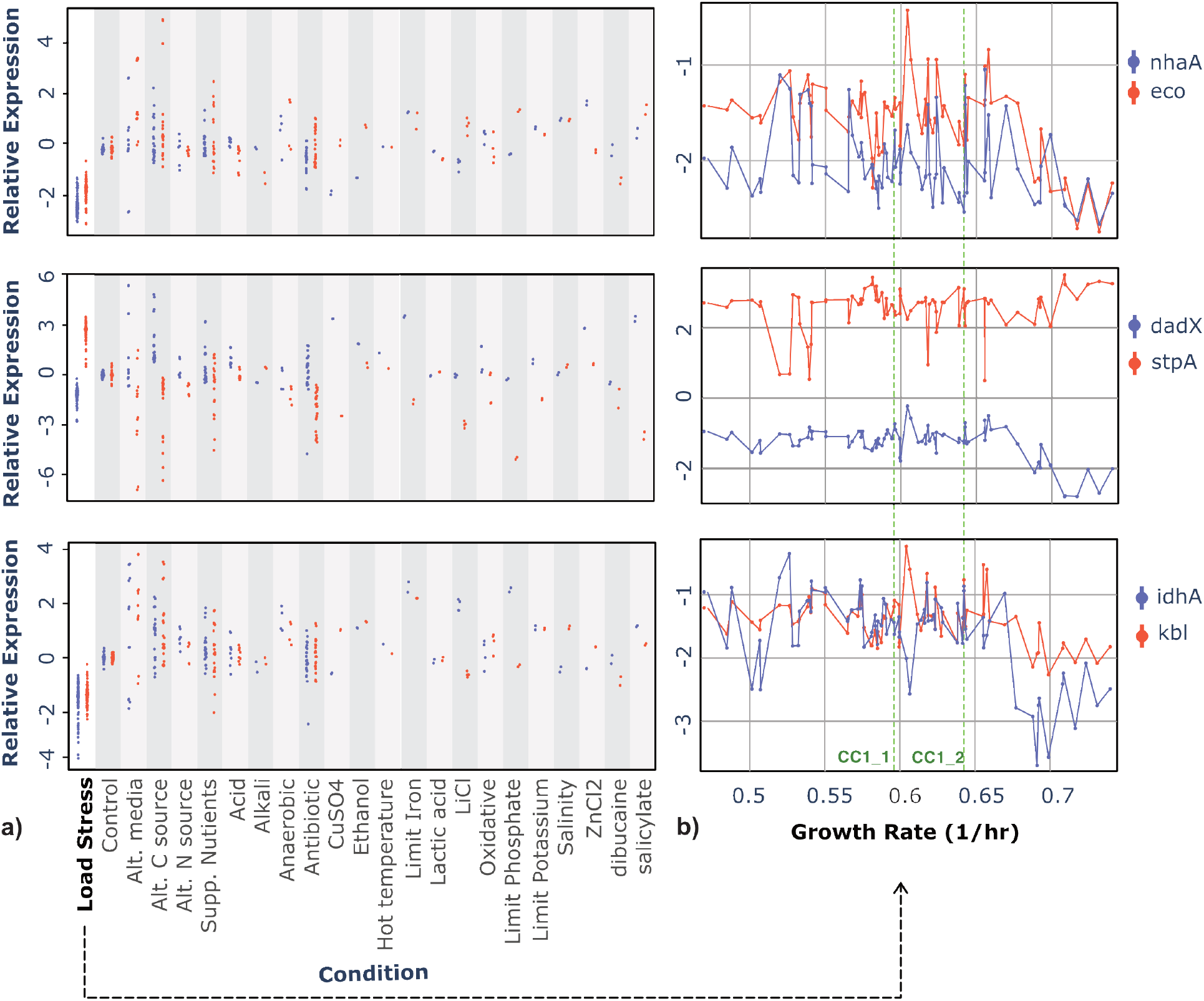
The expression levels of load stress biomarkers and the growth rates of the engineered *E. coli* strains. **a)** The expression levels of biomarker genes are represented as jittered marks in strip plots. Each column shows the gene expression in a different condition, with the load stress in the first column as compared to various other conditions in the rest 22 columns. Two genes in a biomarker pair are discriminated in colour blue and red. **b)** The line plots show the gene expression levels of three biomarker pairs against the cellular growth rates in load stress samples. Two genes in a biomarker pair are discriminated in colour blue and red. Samples inserted with empty plasmid (CC1 1, CC1 2) were marked with green dash lines.

As the load stress can be induced by expressing different synthetic constructs in *E. coli* samples and affecting the cellular growth rate, these samples may suffer from different levels of load stress and reduced growth. We explored the biomarker gene expression in the context of cellular growth rate in 82 load stress samples (Figure 4b). Linear correlation between biomarker expression levels and growth rates were not observed. Surprisingly, we found that among these load stress samples those with higher growth rate (more than 0.68) showed larger scale of upregulation or downregulation of biomarker expression than those with lower growth rate. The pair of samples with empty plasmid inserted (marked by green line in Figure 4b) showed moderate degradation in growth rate and were considered under load stress as well. The interactive version of Figure 4 that enables the hover text with details of each sample dot can be assessed at https://neverbehym.github.io/TBecoli_supp.html.

## 4 Discussion

The expression of synthetic constructs for heterologous protein production in host bacteria can lead to resource and energy competition [32, 33], triggering a load stress response and compromising the performance of the constructsThe goal of this study was to identify the genes that were uniquely responsive to load stress in *E. coli*, so as to enable effective engineering strategies for relieving the cellular load stress and stabilising the heterologous protein production. This was achieved by designing computational methods to exploit a RNA-seq dataset that measured the gene expression profiles of non-induced *E. coli* cells with diverse environmental perturbations and induced *E. coli* cells with a large set of heterologous genes inserted.

The differential expression analysis showed that complex transcriptional adaptions occurred in the context of load stress, which were related to various biological processes such as carbon and Amino Acid Metabolism, Translation, Membrane Homeostasis, Heat Stress and Anti Toxin. We performed an ensemble of feature selection methods to minimise the number of genes required to discriminate load stress state from normal and other stress states, avoiding the selection of some genes over others as biased by certain classifier or feature elimination manner. We found that pairs of genes can already predict load stress samples with accuracy higher than 0.97. Our biomarker gene pairs can be further verified and readily incorporated in the most recent burden feedback system from [20]. The system specificity is expected to improve because the biomarker genes identified in this work were unique to load stress—they can discriminate the load stress from diverse cellular states that might be triggered in other stress conditions. The system sensitivity is also expected to improve because we included 41 different engineered *E. coli* strains in our study as compared to only four different engineered *E. coli* strains in [20].

As this work only studied a single RNA-seq dataset where the engineered *E. coli* strains used same plasmid design, it is preferred to perform validation of our methods on additional synthetic constructs expression from independent samples. Future work also includes the study of the correlation between load sensing gene expression levels and host cell growth rates as well as synthetic construct expression levels.

In this work we provided a quantitative approach to identify pairs of gene signatures that can accurately predict load stress state induced by synthetic construct expression with respect to other cellular states. The biomarker genes identified in this work can be further applied to build a load stress reporting system.

